# Imaging Flow Cytometry reveals a dual role for exopolysaccharides in biofilms: To promote self-adhesion while repelling non-self-community members

**DOI:** 10.1101/2021.09.22.461363

**Authors:** Harsh Maan, Tatyana L. Povolotsky, Ziv Porat, Ilana Kolodkin-Gal

## Abstract

In nature, bacteria are establishing differentiated communities referred to as biofilms. These multicellular communities are held together by self-produced polymers that allow the community members to adhere to the surface as well as to neighbor bacteria. Here, we report that exopolysaccharides prevent *Bacillus subtilis* from co-aggregating with a distantly related bacterium *Bacillus mycoides*, while maintaining their role in promoting self-adhesion and co-adhesion with phylogenetically related bacterium, *Bacillus atrophaeus*. The defensive role of the exopolysaccharides is due to the specific regulation of bacillaene. Single cell analysis of biofilm and free-living bacterial cells using imaging flow cytometry confirmed a specific role for the exopolysaccharides in microbial competition repelling *B. mycoides*. Unlike exopolysaccharides, the matrix protein TasA induced bacillaene but inhibited the expression of the biosynthetic clusters for surfactin, and therefore its overall effect on microbial competition during floating biofilm formation was neutral. Thus, the exopolysaccharides provide a dual fitness advantage for biofilm-forming cells, as it acts to promote co-aggregation of related species, as well as, a secreted cue for chemical interference with non-compatible partners. These results experimentally demonstrate a general assembly principle of complex communities and provides an appealing explanation for how closely related species are favored during community assembly. Furthermore, the differential regulation of surfactin and bacillaene by the extracellular matrix may explain the spatio-temporal gradients of antibiotic production within biofilms.

## Introduction

In terrestrial microenvironments, microbial communities often provide beneficial effects to other organisms, e.g., biocontrol agents grown on the surface of plant roots, thereby preventing the growth of bacterial and fungal pathogens [1].This beneficial effect is often associated with the production of antimicrobial substances and antibiotics [1]. Antibiotic production was shown to be regulated as well as affected by the biological, chemical and physical features of the environment [2-4]. However, the guiding rule for coordinating antibiotic production with the biotic environment remains unknown.

In many natural scenarios, bacteria grow in heterogeneous communities organized into a complex 3D structure, designated as biofilms [5]. The 3D structure of the biofilm was suggested to relieve metabolic stress, by the utilization of channels formed below the ridges and wrinkles within the colony that may facilitate diffusion of fluids, nutrients and oxygen [6-9].

The formation of a biofilm is a developmental process, in which various genetic programs are activated in a specific order in different subpopulations of cells, for the proper establishment of a functional structure [7, 9-14]. This apparent coordination can be explained by the temporally distinct exposure of cell subpopulations to specific microenvironments [13].

To form a functional structure, biofilm-forming cells produce polymers that constitute the extracellular matrix (ECM), where they bind to each other and to the surface. The ECM plays an important role in the resistance and resiliance of the entire biofilm community [15-17]. Although the ability to generate an ECM appears to be a common feature of multicellular bacterial communities, there is remarkable diversity in the means by which these matrices are constructed [18]. The most extensively studied components of biofilm ECMs are carbohydrate-rich polymers (i.e., extracellular polysaccharides or exopolysaccharides (EPS)), proteins, nucleic acids [18, 19] and biogenic minerals [20, 21].

*Bacillus subtilis* is a spore-forming facultative anaerobe and is a commensal that is typically found in the rhizosphere and soil (Errington 1993, Hilbert & Piggot 2004, Reverdy et al 2018). This bacterium often serves as a model organism for beneficial Gram-positive bacteria, and its undomesticated strains form robust and architecturally complex biofilms (Branda et al 2001, Chen et al 2013).

*B. subtilis* is capable of forming various types of biofilms including on solid surfaces (colony), in the air-water interface (pellicles or floating biofilms) and submerged biofilms in domesticated strains. The ECM of *B. subtilis* is characterized by the exopolysaccharides encoded by the *epsA-O* operon and the proteins TasA and BslA [22, 23]. TasA is a functional amyloid and is encoded by the *tapA-sipW-tasA* operon [22, 24, 25]. The <u>b</u>iofilm <u>s</u>urface layer protein <u>A</u> (BslA) is involved in the formation of a water-repellent hydrophobic coat over the biofilm and is critical for pellicle and colony biofilms [26, 27]. All matrix components were expressed and promoted during pellicle formation. Interestingly, all extracellular matrix components were shown to have additional non-structural roles: The protein TasA was shown to regulate motility, extracellular matrix production and the general stress response [28, 29] and exopolysaccharides were shown to activate the master regulator Spo0A [30] which regulates biofilm formation, sporulation and the production of the antibiotic Bacillaene [31-33]. Therefore, the role of the extracellular matrix in microbial competition could be multifactorial due to its independent structural and non-structural roles.

Here, we used imaging flow cytometry to evaluate the roles of all major extracellular matrix components in microbial competition for floating biofilm (pellicle) formation. The population within *B. subtilis* biofilms and root associated communities is heterogeneous [14, 34, 35], a heterogenicity which is affected by the production of the extracellular matrix. To overcome this fundamental property of biofilms, we monitored the effects of the extracellular matrix on the single cell level, relying on imaging flow cytometry. This method combines the power and speed of traditional flow cytometers with the resolution of the microscope. It therefore allows for high rate complex morphometric measurements in a phenotypically defined way [36], and was recently shown to be highly accurate at monitoring genes expressed in subpopulations of *B. subtilis* biofilms [37].

We uncovered that exopolysaccharides, but not the proteinous matrix components, have a dual role in excluding non-self-community members from mixed communities while promoting the self-aggregation and co-aggregation with related species. This role was indirect, and could be attributed to the exopolysaccharides dependent production of bacillaene within floating biofilms.

## Results

### Conditioned medium from *B. subtilis* pellicles is selectively toxic towards potential competitors

To test the effect of secreted products on the competition between related and non-related floating biofilms (pellicles), we collected the conditioned growth medium from *B. subtilis* pellicles and assessed its bioactivity towards three related *Bacillus* species: *Bacillus subtilis* (self), the highly related bacterium *B. atrophaeus*, and the phylogenetically distinct bacterium *B. mycoides. B. subtilis* clade members: *B. subtilis* and *B. atrophaeus* formed pellicles which were enhanced when supplemented with CM from *B. subtilis* (Figure 1A). For *B. mycoides*, no pellicle formation was observed (control), however, growth was observed as cell aggregates were present in suspension.

**Figure 1.**
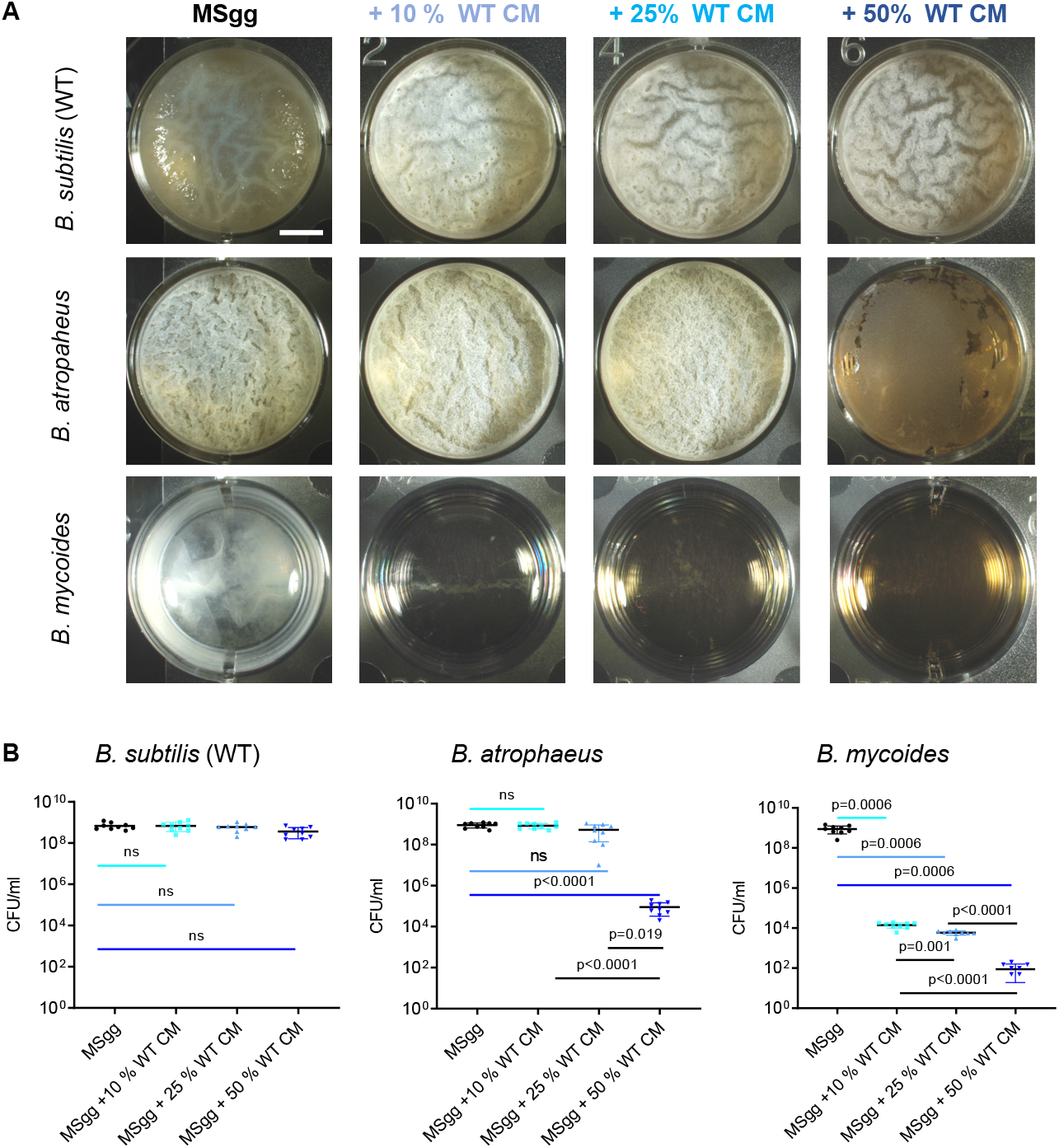
Conditioned medium from *B. subtilis* pellicles is selectively toxic towards potential competitors. **A)** Monitoring pellicle formation and growth of the indicated strains when grown in MSgg medium and MSgg medium supplemented with conditioned medium (CM) from WT *B. subtilis* in increasing concentrations (10%,25% and 50% v/v). Cells were grown at 30°C and images were obtained at 72 h post inoculation and are representative of data presented in B. **B)** The number of CFU obtained when indicated strains were either grown in MSgg medium or MSgg medium supplemented with conditioned medium (CM) from WT *B. subtilis* in increasing concentrations (10%,25% and 50% v/v). Cells were grown at 30°C, and were harvested at 72 h post inoculation. Graphs represent mean ± SD from three independent experiments (n = 9). Statistical analysis was performed using Brown-Forsthye and Welch’s ANOVA, with Dunnett’s T3 multiple comparisons test. P < 0.05 was considered statistically significant. Scale bar = 5mm

A dose-dependent response analysis of all examined species for pellicle formation confirmed a correlation between the relatedness of the strains and their response to the conditioned medium of *B. subtilis* (Figure 1B). *B. subtilis* was fully resistant to its own conditioned medium, *B. atrophaeus* was resistant to low concentrations of, but vulnerable to high concentrations of the conditioned medium, (50%v/v., p value<0.0001) and *B. mycoides* was sensitive to all of the tested concentrations of conditioned medium (p=0.0006) (Figure 1). These results are consistent with our previous work where we reported that the conditioned rich medium of *B. subtilis* pellicles was inert to self, slightly toxic to the related *B. atrophaeus*, and significantly toxic to *B. mycoides* [38].

### The exopolysaccharides positively regulate the selective toxicity of the conditioned medium

To compare the potential non-structural roles of the extracellular matrix components of *B. subtilis* on interspecies competition, we collected the conditioned media of all ECM mutants (Figure 2A) and compared its bioactivity to the conditioned media of the parental strain on recipient cultures. As shown, a deletion of the biosynthetic clusters of the exopolysaccharides (Δeps) significantly reduced the toxicity of the conditioned medium towards *B. mycoides*. Quantifying *B. mycoides* cells grown in the presence of conditioned medium from all donors, confirmed the specific effect of the deletion of the exopolysaccharides resulting in a significant rescue of *B. mycoides*. To test whether this non-structural role of exopolysaccharides was direct, we then supplemented the medium with increasing concentrations of either purified exopolysaccharides (eps) or TasA protein to study if they have any effect on the growth of *B. mycoides*. Interestingly, instead of having any inhibitory effect on the growth, these purified ECM components facilitated the growth of *B. mycoides* (Figure S1 and S2), potentially serving as carbon and nitrogen sources. This suggested that ECM components were not directly responsible for toxicity towards *B. mycoides*. Therefore, we hypothesized that the antagonistic effect of the extracellular matrix and the exopolysaccharides towards the phylogenetically distinct competitor *B. mycoides* is indirect.

**Figure 2.**
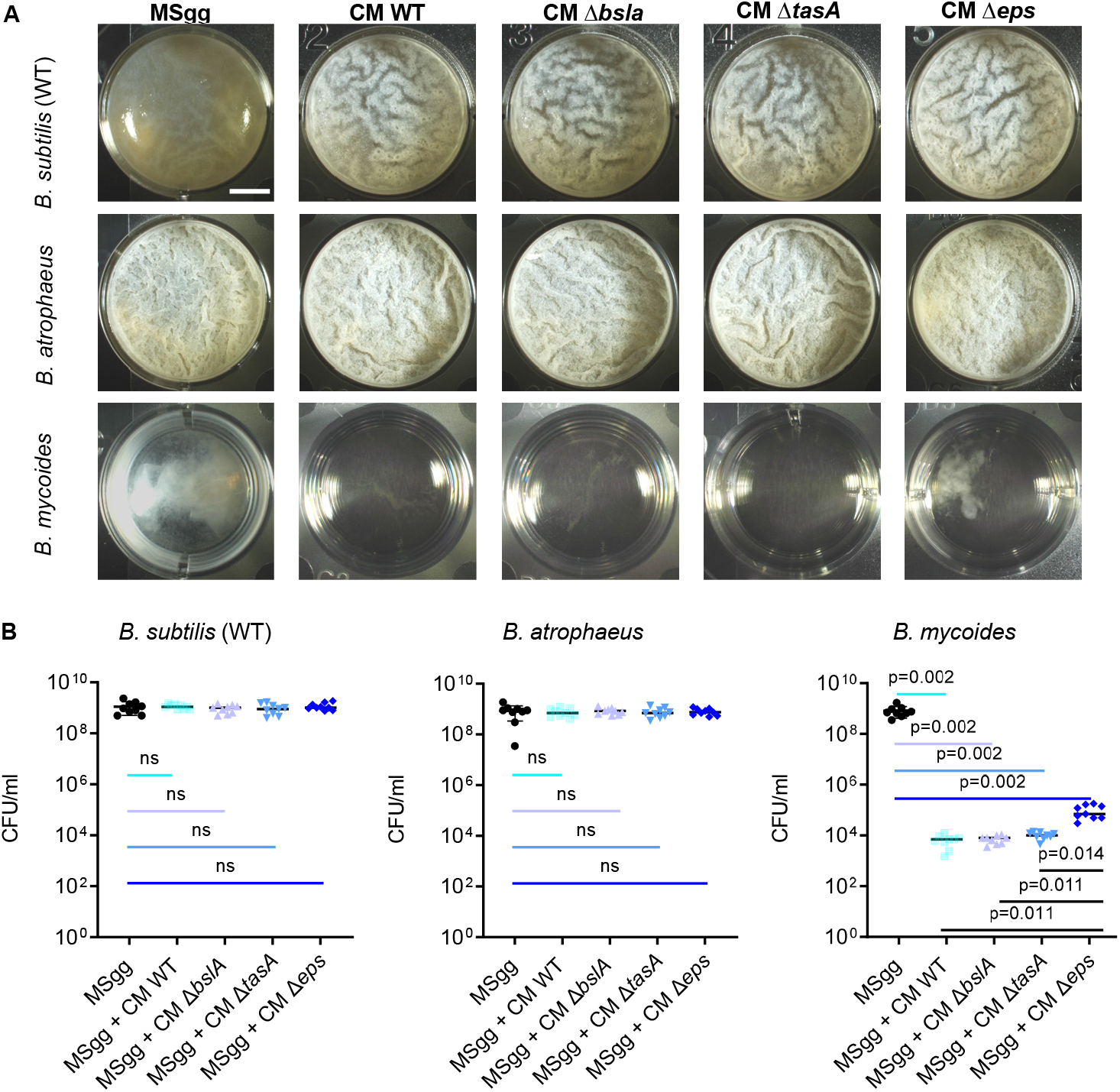
The exopolysaccharides positively regulate the selective toxicity of the conditioned medium. **A)** Monitoring pellicle formation and growth of the indicated strains when grown in MSgg medium and MSgg medium supplemented with conditioned medium (CM) from WT *B. subtilis* and its ECM mutants (10%, v/v). Cells were grown at 30°C and images were obtained at 72 h post inoculation and are representative of data presented in B. **B)** The number of CFU obtained when indicated strains were either grown in MSgg medium or MSgg medium supplemented with conditioned medium (CM) from WT *B. subtilis* and its ECM mutants (10%, v/v). Cells were grown at 30°C, and were harvested at 72 h post inoculation. Graphs represent mean ± SD from three independent experiments (n = 9). Statistical analysis was performed using Brown-Forsthye and Welch’s ANOVA, with Dunnett’s T3 multiple comparisons test. P < 0.05 was considered statistically significant. Scale bar = 5mm

To assess the structural and non-structural roles of exopolysaccharides and proteinous matrix during interspecies competitions, we competed *B. subtilis* and its matrix mutants versus itself, *B. atrophaeus* (mostly resistant to the conditioned media) and *B. mycoides* (mostly sensitive to the conditioned media) for pellicle formation. Pellicle formation is a system which is less prone to diffusion limitations as compared with semi-solid agar surfaces, and therefore allows us to better quantify the response of the different competitors to secreted products. For strains capable of forming a pellicle, we compared the presence of the different competitors within the pellicle and within the suspension separately. To get accurate results we relied on imaging flow cytometry, a technique which allows us to image the cells and to fully distinguish between and accurately quantify constitutively expressing *B. subtilis* P_*hyperspank*_*-gfp* (608) or its ECM mutants vs their competitor’s: WT *B. subtilis, B. atrophaeus* and *B. mycoides*.

As shown in Figure 3A and 3B, *B. subtilis* was capable of forming pellicles in an exopolysaccharide-dependent manner with itself and with *B. atrophaeus* as well as to co-aggregate with these strains. During competition with *B. mycoides* no significant differences were observed in pellicle formation of *B. subtilis* WT and its ECM mutants (Figure S5) regardless of the presence of the extracellular matrix, indicating a barrier for joint pellicle formation. However, the presence of free living *B. mycoides* cells in suspension was detected and these free living cells co-existed significantly better with *B. subtilis* strain lacking the exopolysaccharide biosynthetic operon (Δ*eps*) (p=0.022) than with the WT (Figures 3C and D).

**Figure 3.**
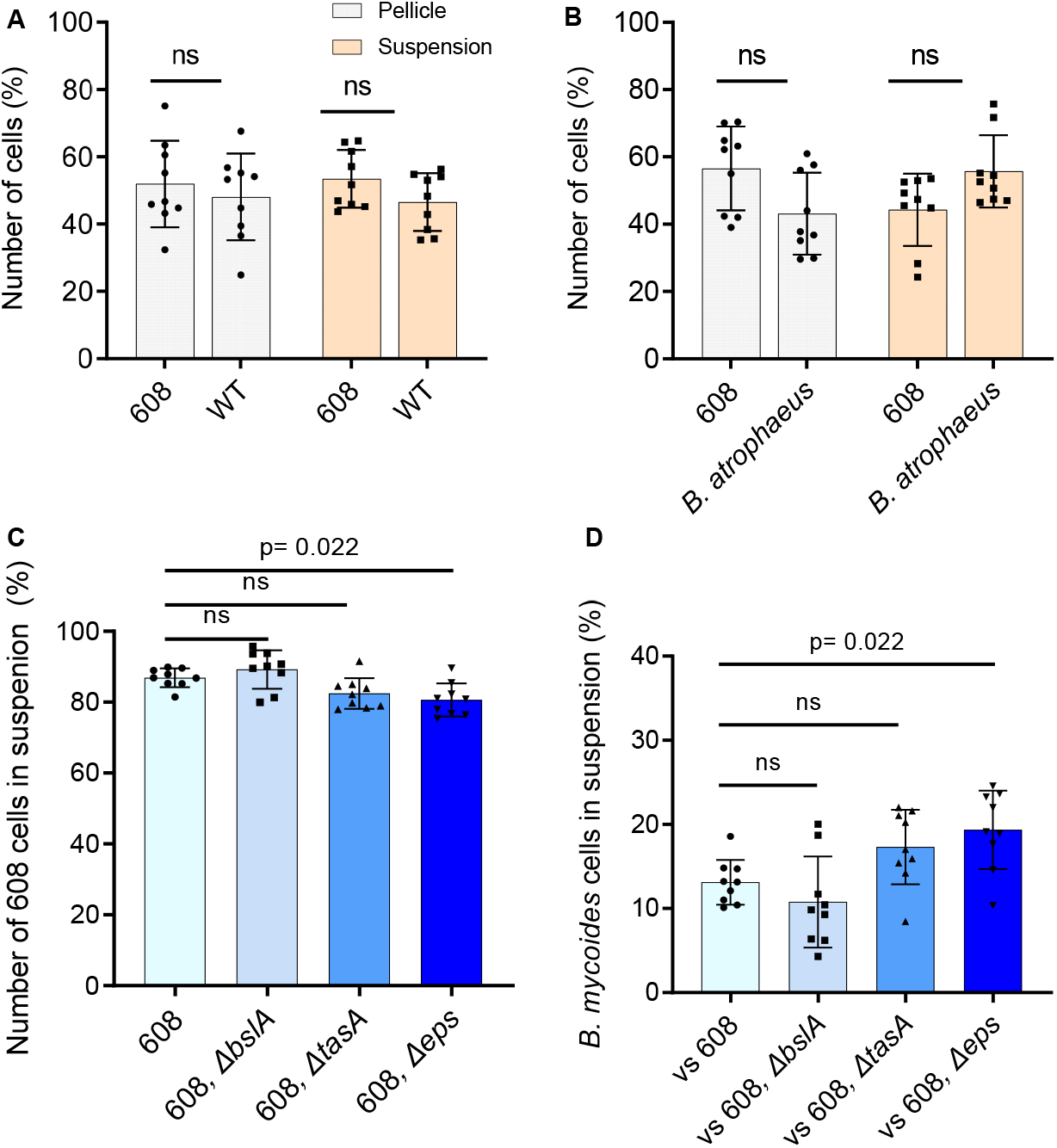
Exopolysaccharides prevent growth of free-living cells of the sensitive competitor *B. mycoides*. Quantifying the number of cells present in the pellicle/suspension of each competition assay using imaging flow cytometry. **A)** A competition assay was set up between WT *B. subtilis* stains harboring P_*hyerspank*_-*gfp* (608), and WT *B. subtilis* in MSgg medium. Quantification of number of cells of each strain both in pellicle and suspension was performed using imaging flow cytometry. **B)** A competition assay was set up between WT *B. subtilis* stains harboring P_*hyerspank*_-*gfp* (608), and *B. atrophaeus* in MSgg medium. Quantification of number of cells of each strain both in pellicle and suspension was performed using imaging flow cytometry. **C)** A competition assay was set up between WT *B. subtilis* stains harboring P_*hyerspank*_-*gfp* (608), 608, Δ*tasA*, 608 Δ*bslA* and 608 Δ*eps* mutants vs *B. mycoides*, in MSgg medium. From the suspension culture, number of 608 and its deletions mutants cells, in competition against *B. mycoides* cells were quantified using imaging flow cytometry. **D)** A competition assay was set up between WT *B. subtilis* stains harboring P_*hyerspank*_-*gfp* (608), 608 Δ*tasA*, 608 Δ*bslA* and 608 Δ*eps* mutants vs *B. mycoides*, in MSgg medium. From the suspension culture, number of *B. mycoides* cells in competition against 608 and its deletion mutant cells were quantified using imaging flow cytometry. Data were collected 72 h post inoculation, and 100,000 cells were counted. Graphs represent mean ± SD from three independent experiments (n=9). The data were analyzed using IDEAS 6.3. For A-B, statistical analysis was performed using two-way ANOVA followed by Dunnett’s multiple comparison test. For B-C, statistical analysis was performed Brown-Forsthye and Welch’s ANOVA, with Dunnett’s T3 multiple comparisons test. P < 0.05 was considered statistically significant.

### Exopolysaccharides regulate the antibiotic bacillaene to eliminate sensitive competitors

Recently, we found that two antibiotics: the NRP (non-ribosomal peptide) surfactin and the polyketide bacillaene play an important role in the elimination of phylogenetically distinct competitors over semi-solid surfaces and in shaking culture [38]. As we found that the role of exopolysaccharides (Δ*eps*) during competition is largely indirect, and was pronounced in the free-living cells present in suspension, we tested whether the exopolysaccharides are capable to regulate surfactin and bacillaene production. The expression of operons encoding surfactin and bacillaene biosynthetic clusters tagged to a luciferase reporter [38] were compared for ECM mutants. Both exopolysaccharides and TasA were required for optimal bacillaene expression (Figure 4A); however, a deletion of TasA induced surfactin production (Figure S3). These results could explain why deletion of Δ*eps* has a more pronounced effect on the toxicity towards *B. mycoides*, compared with Δ*tasA*.

**Figure 4.**
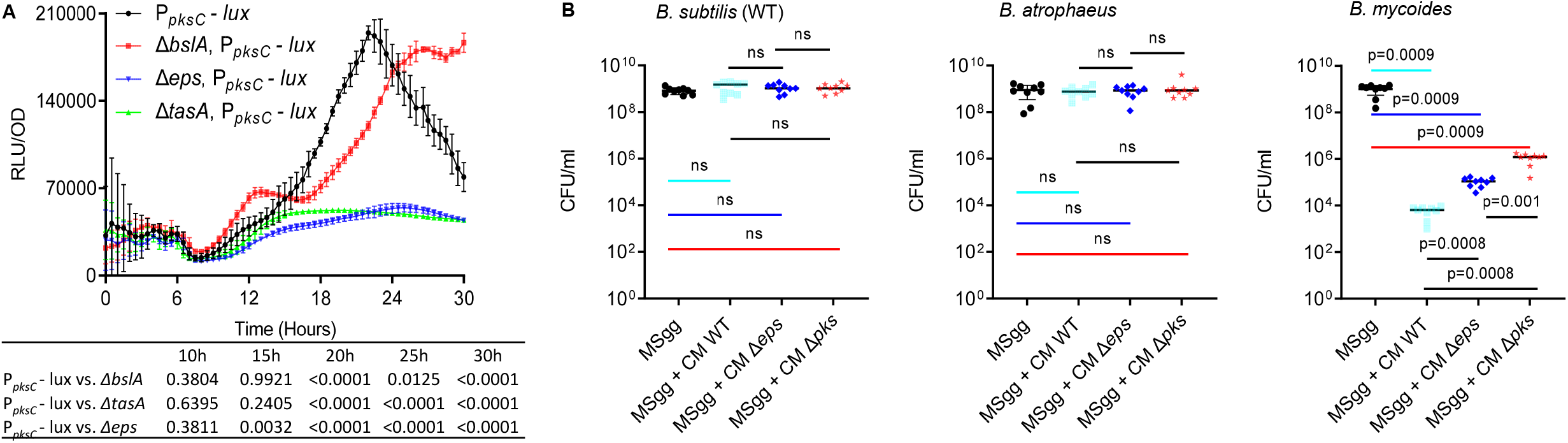
Exopolysaccharides and the antibiotic Bacillaene are essential to eliminate sensitive competitors. **A)** Analysis of the luciferase activity in a WT *B. subtilis* strain harboring P_*pksC*_*-lux* (bacillaene) reporter, and its indicated ECM deletion mutants. Luminescence was monitored in MSgg medium, at 30°C for 30 h. Graphs represent mean ± SD from three independent experiments (n = 9). Statistical analysis was performed using two-way ANOVA followed by Dunnett’s multiple comparison test. P < 0.05 was considered statistically significant. P values at different time points are shown in the table. Luciferase activity was normalized to avoid artifacts related to differential cell numbers as RLU/OD. **B)** The number of CFU obtained when indicated strains were either grown in MSgg medium or MSgg medium supplemented with conditioned medium (CM) from WT *B. subtilis* and its Δ*eps* and Δ*pks* mutants (10%, v/v). Cells were grown at 30°C, and were harvested at 72 h post inoculation. Graphs represent mean ± SD from three independent experiments (n = 9). Statistical analysis was performed using Brown-Forsthye and Welch’s ANOVA, with Dunnett’s T3 multiple comparisons test. P < 0.05 was considered statistically significant.

To test whether the deletion of bacillaene biosynthetic clusters indeed mimics the effect of deletion of the exopolysaccharides, we examined the effect of a *pks* mutant (deficient in bacillaene biosynthesis) on the bioactivity of the conditioned medium. Accordingly, the deletion of *pks* significantly reduced the toxicity of the conditioned medium towards *B. mycoides* (Figure 4B, S5).

To confirm that matrix exopolysaccharides act independently of their structural role to induce bacillaene, we quantified the expression from the promoter of *pks* tagged to a GFP reporter using imaging flow cytometry in the parental strains, and its *eps* mutant. As shown (Figure 5) the intensity of the expression from *pks* promoter and the amount of positively expressed cells were significantly reduced in the exopolysaccharides mutant (Δ*eps*) throughout static growth. For WT *B. subtilis* cells growing in suspension, a reduced expression was observed as compared to the cells growing within the pellicle, however, cells growing in suspension also demonstrated a higher expression from *pks* promoter compared with the exopolysaccharides mutant. These results confirm a non-structural role for exopolysaccharides in inducing bacillaene, which contributes to the favorable outcome of interspecies interactions in biofilms.

**Figure 5.**
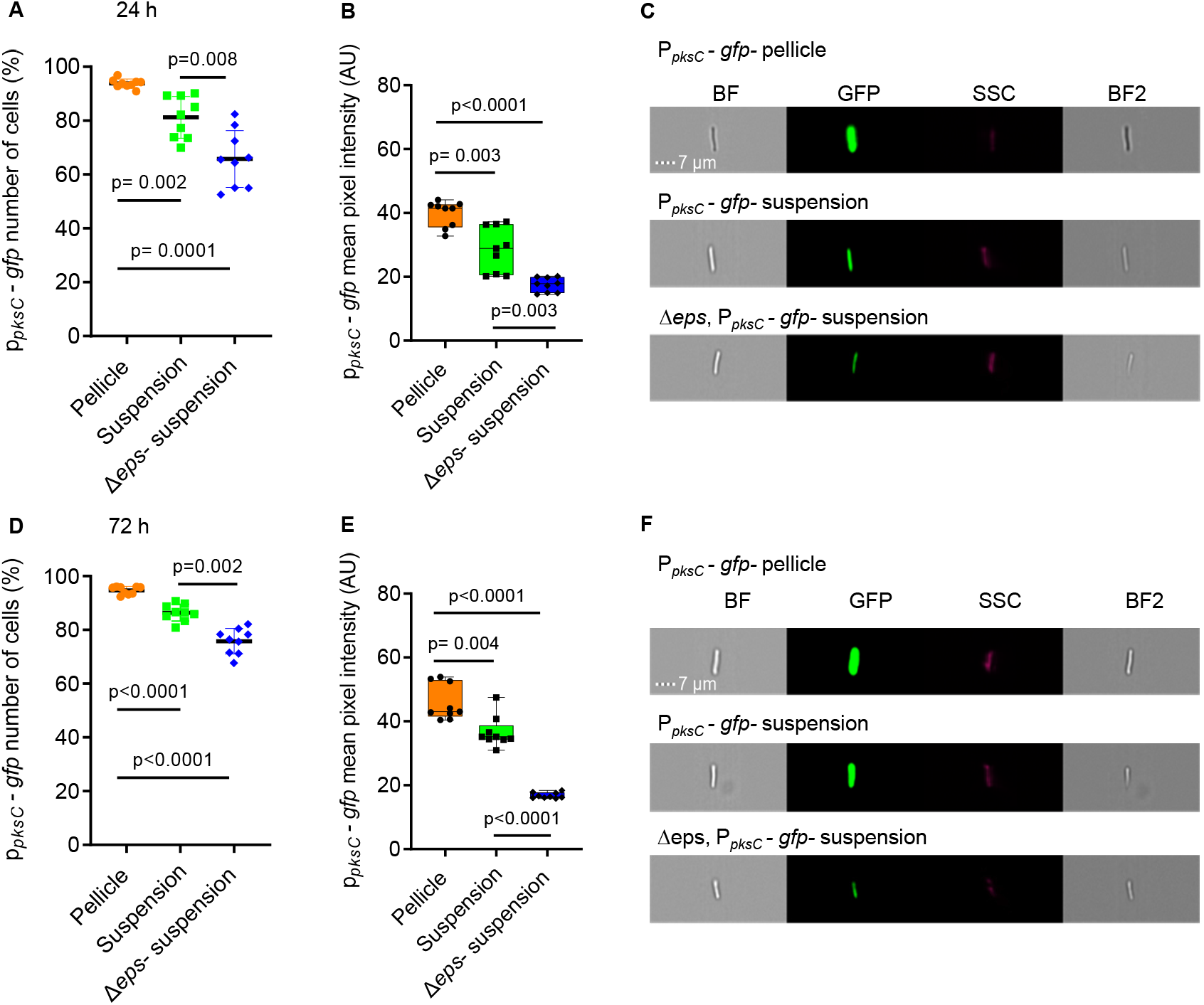
Exopolysaccharides induce bacillaene expression. Monitoring the expression of bacillaene in WT *B. subtilis* strain and its Δ*eps* mutant harboring P_*pks*_-*gfp* reporter. The bacillaene expression was measured using imaging flow cytometry in both pellicle and suspension in for WT B. subtilis P_*pks*_-*gfp*, and only in suspension for Δ*eps, B. subtilis* P_*pks*_-*gfp* mutant, due to its inability to form pellicle. **A)** Showing positively expressing fluorescent populations (%) at 24 h. **B)** Mean pixel intensity of the fluorescent positive populations as in A. **C)** Representative bright-field and fluorescent images related to expression of reporters in A, scale bar = 7µm. **D)** Showing positively expressing fluorescent populations (%) at 72 h. **E)** Mean pixel intensity of the fluorescent positive populations as in A. **F)** Representative bright-field and fluorescent images related to expression of reporters in F, scale bar = 7µm. Graphs A-D represent mean ± SD from three independent experiments (n = 9). Graphs B-E represent box and whiskers plots showing median and interquartile range together with maximum and minimum values and outlier points from three independent experiments (n = 9). Statistical analysis was performed using Brown-Forsthye and Welch’s ANOVA, with Dunnett’s T3 multiple comparisons test. P < 0.05 was considered statistically significant.

## Discussion

Bacteria in nature are most often found in the form of multicellular aggregates commonly referred to as biofilms [39, 40]. When compared to the planktonic (free-living) state, cells in biofilms are more protected from environmental insults, including sterilizing agents, antibiotics, and the immune system. Biofilms enable bacteria to attach more firmly to their hosts and better access to nutrients [41-47].

The self-produced extracellular matrix (ECM) surrounds and protects the cells, and makes them adhere to each other or to a surface [19]. This feature makes bacterial biofilms an especially appealing system in which to study multicellular development, with exopolysaccharides, carbohydrate rich polymers being the most studied and wide spread ECM component [18, 48].

Various genetic analyzes have provided strong evidence that biofilm exopolysaccharides play a fundamental structural role in different bacterial species, impact bacterial virulence, and promote capsule formation [49-54]. The biofilm of *B. subtilis* contains several exopolysaccharide polymers, produced by the *epsA-O* operon, and composed of glucose, galactose and N-acetyl-galactosamine [55, 56]. Colonies of mutants in the *epsA-O* operon, and specifically the glycosyltransferase gene *epsH*, lack the exopolysaccharide component of the ECM and are featureless, as opposed to the wrinkled wild type colony [32].

Here we report that in addition to their structural role, manifested by their necessity for *B. subtilis* to generate pellicles (floating biofilms) with itself [57] and with a related bacterium, *B. atrophaeus* (Figure 1), the exopolysaccharides act to induce the production of bacillaene, a polyketide antibiotic [33, 58-60] which was recently shown to be essential for the elimination of phylogenetically distinct *Bacillus s*pecies [38, 61]. Our results indicate that this non-structural role of the exopolysaccharides is manifested by a capacity of *B. subtilis* to repel the phylogenetically distinct competitor, *B. mycoides* from the pellicle and its surrounding media.

Interestingly, the protein matrix component TasA induced Bacillaene while repressing the production of the surfactant and antibiotic surfactin [62, 63], also involved in horizontal gene transfer [64]. This result may explain the heterogeneity in gene expression of antibiotics within the biofilm [38], and predicts that their local expression may also reflect the local concentration of the extracellular matrix components.

So far we could observe two global signals for inducing antibiotic production in *B. subtilis*: peptidoglycan [38] and plant secretions [37] that induced both surfactin and bacillaene. In contrast, ECM driven regulation of antibiotic production is highly specific: TasA represses surfactin and induces bacillaene, and exopolysaccharides trigger the production of bacillaene but to some extent surfactin expression, indicating an overall fine-tuning of antibiotic production.

One advantage of differential ECM-driven regulation is a potential capacity to control local expression within the complicated biofilm structure [18, 65].

When considering a developmental model for biofilm formation, it is tempting to speculate that the bacterial ECM is involved in regulation of genetic programs in designated subpopulation of cells in the biofilm [18]. It has been evident that in multicellular eukaryotes, the production of communication factors depends on cell– ECM interactions [66]. In this work, we describe a similar role for the exopolysaccharides in changing the decision-making processes and the competitiveness of the bacterial biofilm-forming cells. Furthermore, we demonstrate how this ECM-derived cue maintains an essential subset of antibiotic producing cells within the biofilm population.

## Materials and methods

### Strains and Media

All strains used in this study are listed in Supplementary Table 1.

For cloning purposes, selective media was prepared using LB broth or LB-agar using appropriate antibiotics at the following concentrations, 10 μg/ml chloramphenicol (Amersco), 10 µg/ml, 10 μg/ml spectinomycin (Tivan biotech), 10 µg/ml kanamycin (A.G scientific), 1 µg/ml erythromycin (Amresco) + 25 µg/ml lincomycin (Sigma-Aldrich) .MSgg was prepared as described previously [32]. Briefly, 5 mM potassium phosphate, 100 mM MOPS (pH 7), 2 mM MgCl2, 50 μM MnCl2, 50 μM FeCl3, 700 μM CaCl2, 1 μM ZnCl2, 2 μM thiamine, 0.5% glycerol, 0.5% glutamate, Threonine (50 μg ml−1), Tryptophan (50 μg ml−1), and PhenylAlanine (50 μg ml−1).

### Strains construction

All strains are reported in table S1 and were generated using double homologous recombination into neutral integration sites. Briefly, DNA was extracted from the indicated deletion stains using Wizard Genomic DNA Purification Kit (Promega) and transformed into the genome of indicated *B. subtilis* reporter strains induced for natural competence [67]. All mutant strains were confirmed for an accurate integration by PCR.

### Conditioned medium from pellicle culture preparation

Conditioned medium (CM) refers to filtered supernatant of the culture media: Cells from indicated strains were grown to a mid-logarithmic phase of growth (OD=0.6-0.8) and were diluted 1:1000 and suspended in MSgg medium in a 12 well microtiter plate (Corning). Cells were incubated at 30°C for 72 h at the dark. (Brunswick™ Innova® 42). After incubation, the formed pellicle was scraped off, and cells were removed by a centrifugation. The CM was filtered using a 0.22μm filter (Corning). For each independent experiment, a fresh CM was collected.

### Pellicle assays

Cells from a single colony isolated on LB plates were grown to mid-logarithmic phase in 2-ml of LB (∼3 hours at 37°C). Cells were diluted 1:1000 and were added to the 2 mL liquid MSgg in 24 well plates (Thermo Scientific). The cultures were then grown at 30°C for 72 h in the dark. The pellicles were imaged at 72 h. Cells were either grown in the presence or absence of CM as indicated in each corresponding figure legends. The pellicles were imaged using stereomicroscope (Zeiss), using Objective Plain 0.5× FWD 134 mm lens at 10x magnification. Captured images were processed using Zen software (Zeiss).

For CFU count, pellicles were harvested at 72 h by scrapping off the entire pellicle plus pipetting the entire underlying cell suspension, and suspending in PBS (phosphate-buffered saline). This was followed by sonication using a BRANSON digital sonicator, at an amplitude of 10% and a pulse of 5 sec to separate the cells without compromising their viability. 200 µl of sonicated cells suspension was transferred to a Griener 90 well plate (Sigma-Aldrich) and a serial dilution ranging from 10^−1^ to 10^−7^ was performed in PBS. From the dilutions 20 µl of sample was transferred on a LB plate, incubated at 30 °C overnight and then counted.

### Growth Measurements

Cells were grown from a single colony isolated over LB plates to a mid-logarithmic phase of growth. Cells were then grown in 300 μl of MSgg medium in a 96-well microplate (Thermo Scientific), with agitation, at 30°C for 30 h, in a microplate reader (Synergy 2; BioTek, Winooski, VT, USA), and the optical density at 600 nm was measured every 30 min. Cells were either grown in the presence or absence of exopolysaccharides or TasA purified fractions as indicated in each corresponding figures legends.

### Luminescence Analysis

Strains carrying the indicated luminescence reporters were grown from a single colony isolated over LB plates to a mid-logarithmic phase of growth. Cells were then grown in 300 μl MSgg medium, in a 96 well plate with white opaque walls and clear tissue culture treated flat bottoms (Corning). Measurements were performed every 30 min at 30°C for 30 h, using a microplate reader (Synergy 2; BioTek, Winooski, VT, USA). Luciferase activity was calculated as RLU/OD, to avoid artefacts related to the normalization of luminescence intensity to the population size.

### TasA Protein

TasA was expressed and purified as described in [29] and was a kind gift from the Romero lab

### EPS extraction

EPS was extracted from *B. subtilis* pellicles grown for 72 h at 30°C in MSgg medium, as described in pellicle assays. In total 15 pellicle colonies were scrapped and suspended in phosphate-buffered saline (137 mM NaCl, 2.7 mM KCl, 10 mM Na_2_HPO_4_, 1.8 mM KH_2_PO_4_), mildly sonicated, and were centrifuged to remove the cells. The supernatant was collected and mixed with five volumes of ice-cold isopropanol and incubated overnight at 4°C. Samples were then centrifuged at 7,500 × g for 10 min at 4°C. The obtained pellets were suspended in a digestion mix (0.1 M MgCl_2_, 0.1 mg/ml of DNase, and 0.1 mg/ml of RNase), and were incubated for 4 h at 37°C. Following incubation samples were extracted twice with phenol-chloroform. The aquatic fraction was dialyzed for 48 h with Slide-A-Lyzer dialysis cassettes by Thermo Fisher, with a 3,500 molecular weight cut-off, against distilled (d)H_2_O. Samples were lyophilized and lyophilized fraction was dissolved in 500 μl of dH_2_O. The fractions were stored at −80 °C for further use.

### Imaging Flow Cytometry

Indicated strains from the corresponding figure legends were grown in MSgg medium as described in pellicle assays. For flow cytometry analysis pellicle from the pellicle forming strains was gently scrapped off and separated from its suspension and pellicles were suspended in PBS. Both the pellicles and their suspension counterparts were then sonicated using a BRANSON digital sonicator, at an amplitude of 10% and a pulse of 5 sec to separate the cells without compromising their viability. These sonicated samples were then subjected to Imaging Flow Cytometry.

Data was acquired by ImageStream^X^ Mark II (AMNIS, part of Luminex corp., Austin Tx) using a 60X lens (NA=0.9). For GFP excitation lasers intensity was set at 488nm (200mW), and 785nm (5mW) for side scatter measurement. Acquired bacterial cells were gated according to their area (in square microns) and side scatter, during this acquisition, calibration beads which run in the instrument along with the sample were excluded from the gate. Wild type *B. subtilis* was used as a negative control, to separate GFP negative from true GFP positive cells. For each sample, 100,000 events were collected. The acquired data were analysed using IDEAS 6.3 (AMNIS). Every single event represented bacteria that were selected according to their area (in square microns) and aspect ratio (width divided by the length of a best-fit ellipse). Focused events were selected by the Gradient RMS and Contrast features (measures the sharpness quality of an image by detecting large changes of pixel values in the image). GFP positive cells were selected using the Intensity (the sum of the background subtracted pixel values within the image) and Max Pixel values (the largest value of the background-subtracted pixels) of the GFP channel (Ch02). GFP expression was quantified using the Mean Pixel feature (the mean of the background-subtracted pixels contained in the input mask).

### Statistical analysis

All experiments were performed three separate and independent times in triplicates. Datasets were compared using a standard Two-way ANOVA, followed by Tukey’s multiple comparison post hoc testing. For data sets with unequal variances, multiple comparison using Brown-Forsythe and Welch ANOVA tests with Dunnett’s T3 multiple comparison test in order to correct for groups with significantly unequal variances. Error bars represented ± SD, unless stated otherwise. p < 0.05 was considered statistically significant.

Statistical analyses were performed with GraphPad Prism 9.0 (GraphPad Software, Inc., San Diego, CA).

## Supporting Information

Supporting Figures (S1-S5)

Supporting Table (S1)

Supporting References

**Figure S1.**
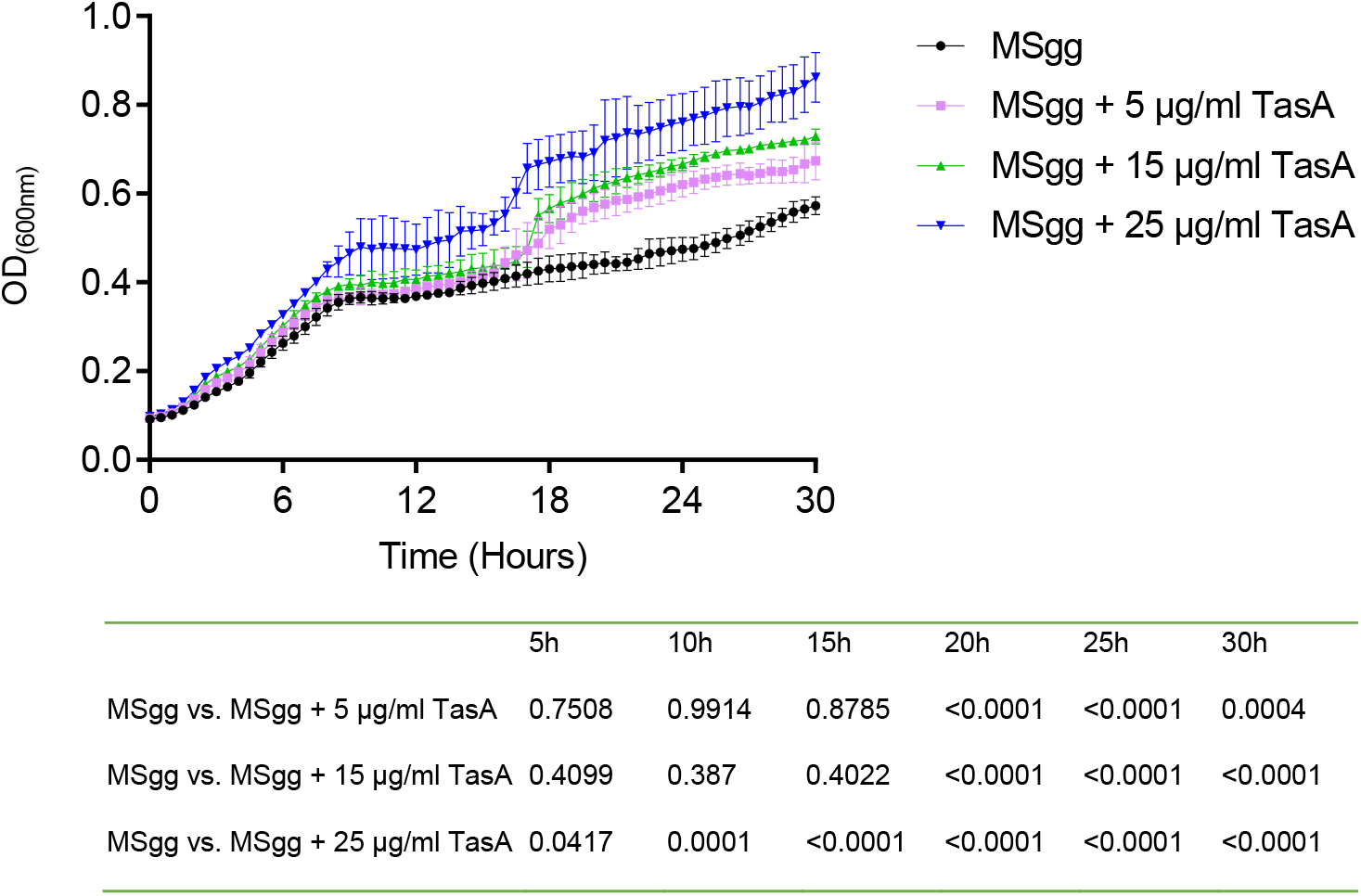
Monitoring growth of *B. mycoides* when supplemented with purified TasA protein. Growth of *B. mycoides* was monitored in MSgg medium and MSgg medium supplemented with the purified TasA protein in increasing concentrations. An increase in the growth was observed in the presence of purified tasA protein. Graph represent the mean ± SD from three independent experiments (n = 9). Statistical analysis was performed using two-way ANOVA followed by Tukey’s multiple comparison post hoc testing. P < 0.05 was considered

**Figure S2:**
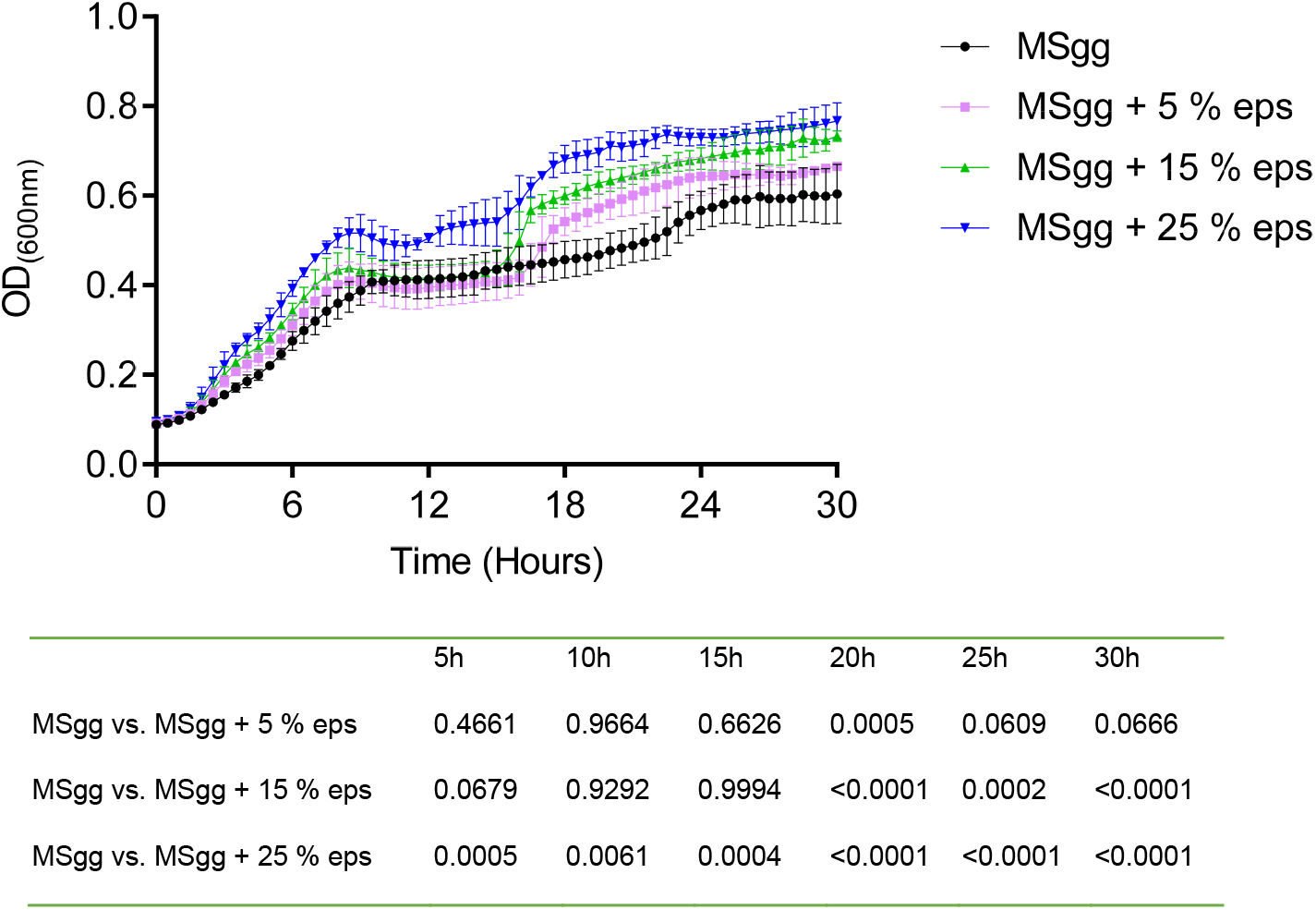
Monitoring growth of B. mycoides when supplemented with purified eps. Growth of *B. mycoides* was monitored in MSgg medium and MSgg medium supplemented with the purified eps in increasing concentrations. An increase in the growth was observed in the presence of purified eps. Graph represent the mean ± SD from three independent experiments (n = 9). Statistical analysis was performed using two-way ANOVA followed by Tukey’s multiple comparison post hoc testing. P < 0.05 was considered statistically significant. P values at

**Figure S3:**
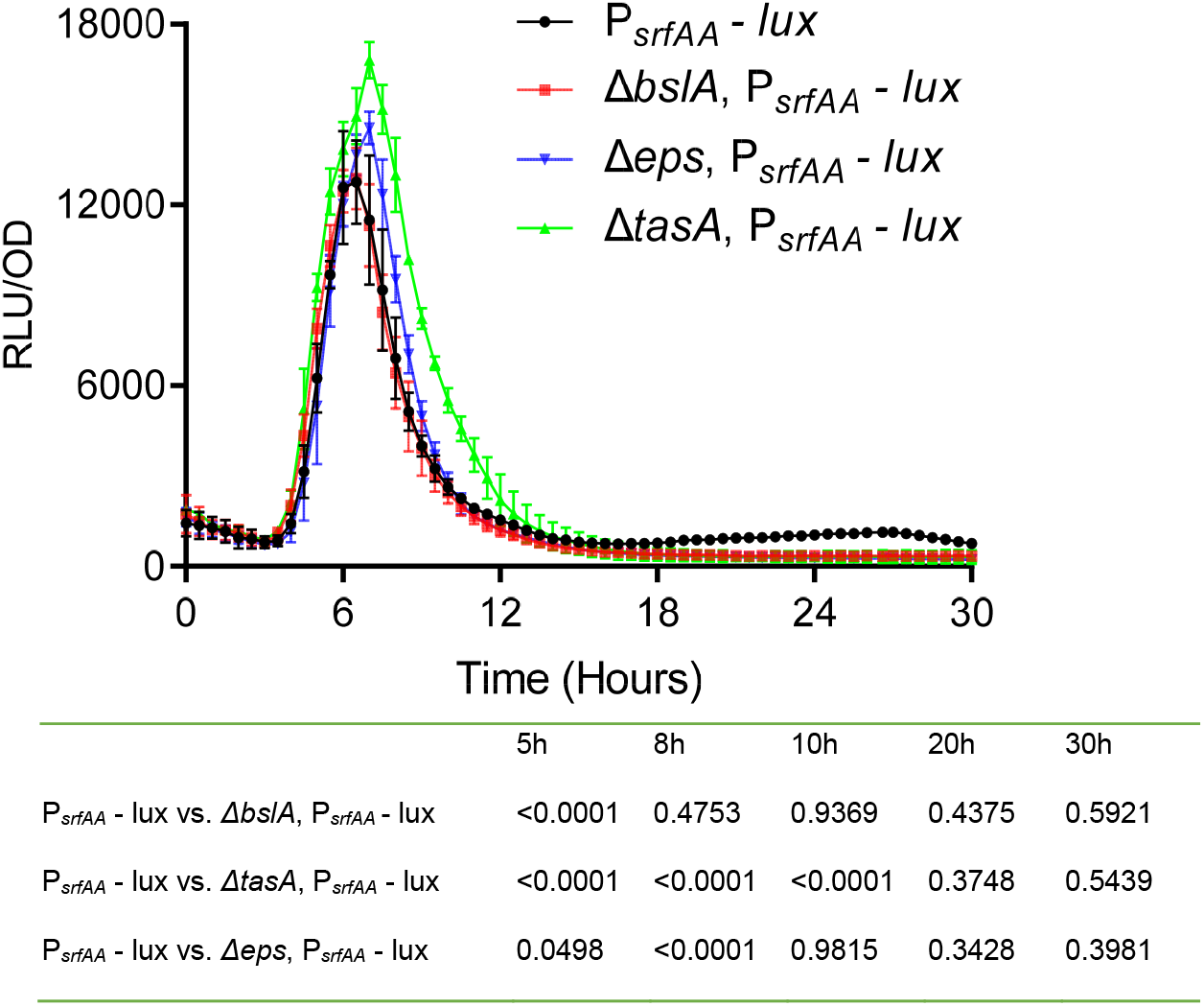
Analysis of the luciferase activity in a WT *B. subtilis* strain harboring P_*srfAA*_*-lux* (surfactin) reporter, and its indicated ECM deletion mutants. Luminescence was monitored in MSgg medium, at 30°C for 30 h. Graphs represent mean ± SD from three independent experiments (n = 9). Statistical analysis was performed using two-way ANOVA followed by Dunnett’s multiple comparison test. P < 0.05 was considered statistically significant. P values at different time points are shown in the table. Luciferase activity was normalized to avoid artifacts related to differential cell numbers as RLU/OD.

**Figure S4:**
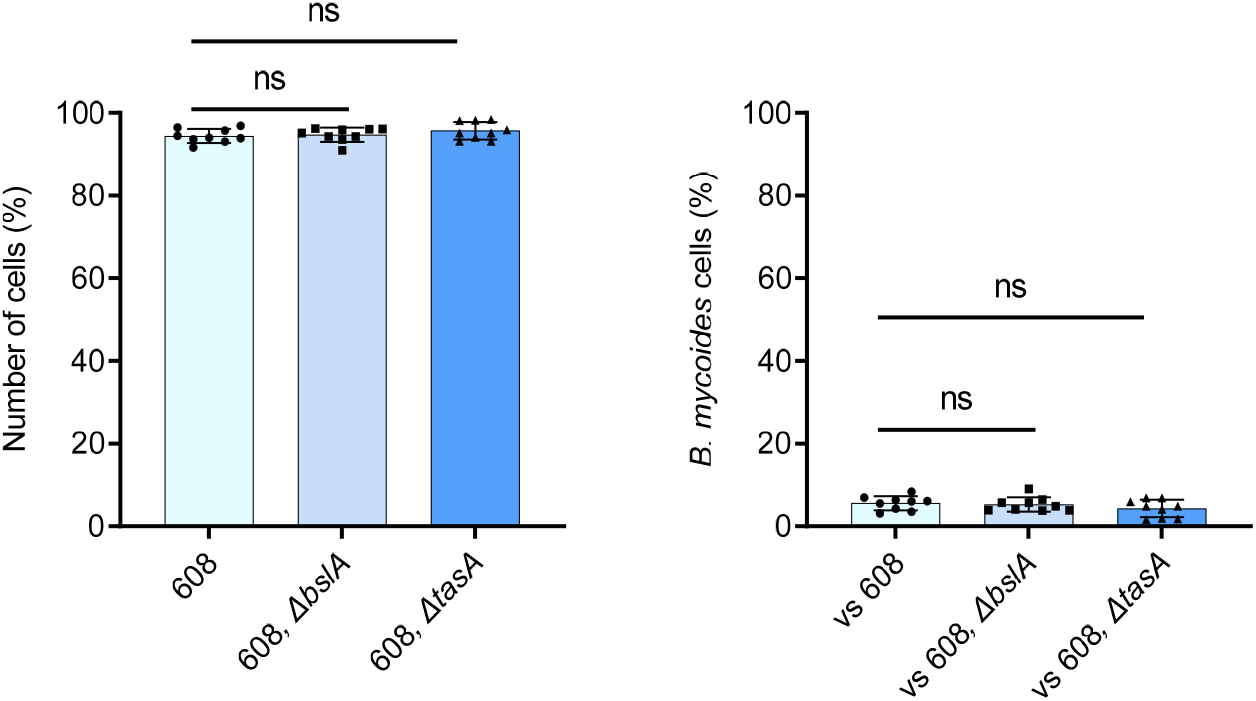
Quantifying the number of cells present in the pellicles of each competition assay using imaging flow cytometry. A competition assay was set up between WT *B. subtilis* stains harboring P _*hyerspank*_-*gfp* (608), 608, Δ*tasA* and 608, Δ*bslA* mutants vs *B. mycoides*, in MSgg medium. As Δ*eps* mutant cannot form pellicles, hence it was not considered for this competition assay. **A)** Quantifying the number of 608 and its deletions mutants cells, in competition against *B. mycoides* cells, using imaging flow cytometry. **B)** Quantifying the number of *B. mycoides* cells in competition against 608 and its deletions mutants using imaging flow cytometry. Data were collected 72 h post inoculation, and 100,000 cells were counted. Graphs represent mean ± SD from three independent experiments (n=9). The data were analyzed using IDEAS 6.3. Statistics analysis was performed using Brown-Forsthye and Welch’s ANOVA, with Dunnett’s T3 multiple comparisons test. P < 0.05 was considered statistically significant.

**Figure S5.**
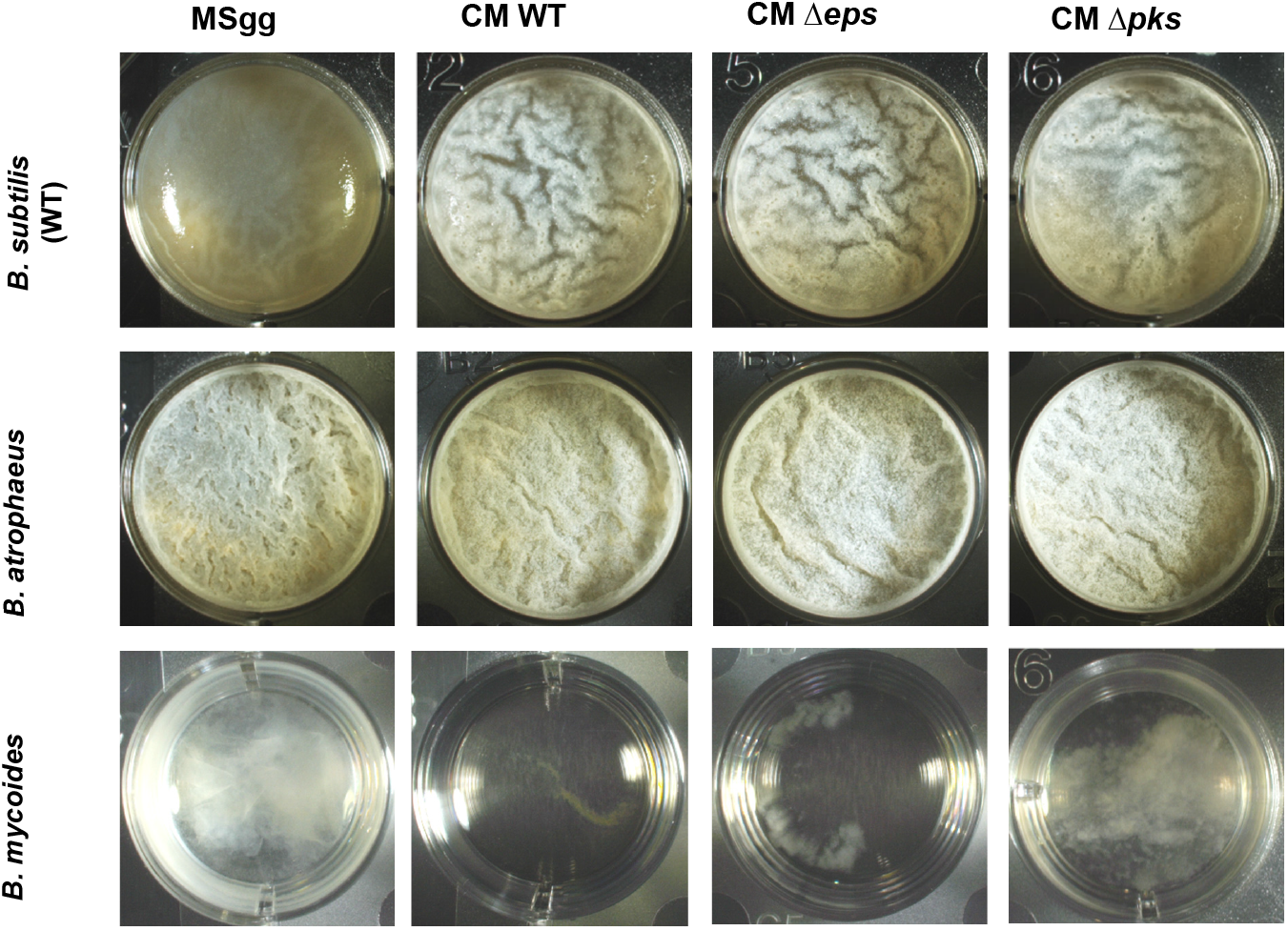
The bioactivity of the CM on pellicle formation. Monitoring pellicle formation and growth of the indicated strains in MSgg medium supplemented with conditioned medium (CM) of WT *B. subtilis* and its Δ*eps* and Δ*pks* mutants (10%, v/v). Cells were grown at 30°C and images were obtained at 72 h post inoculation, and are representative of data shown in Figure 4B.

**Supplementary Table.**
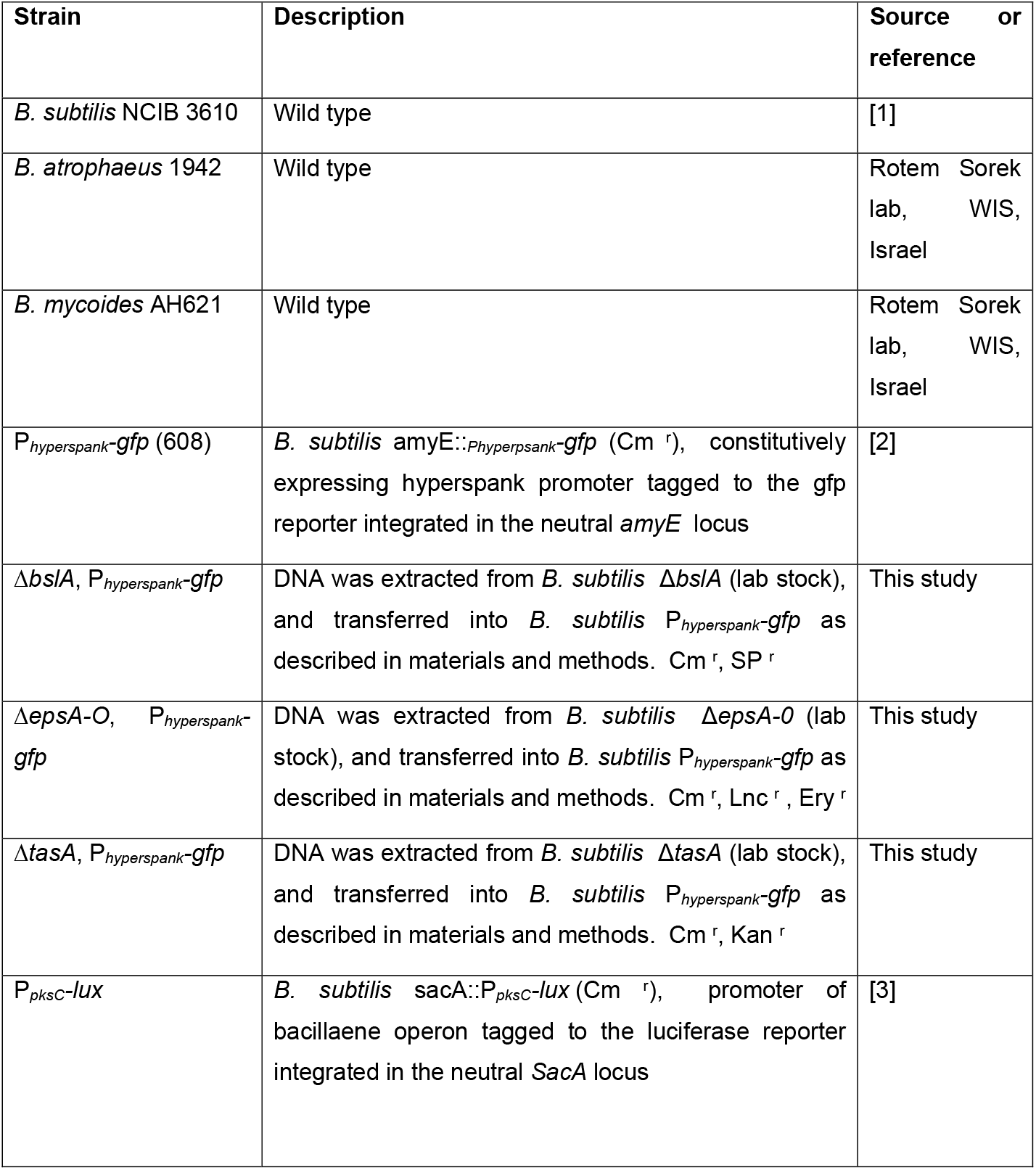
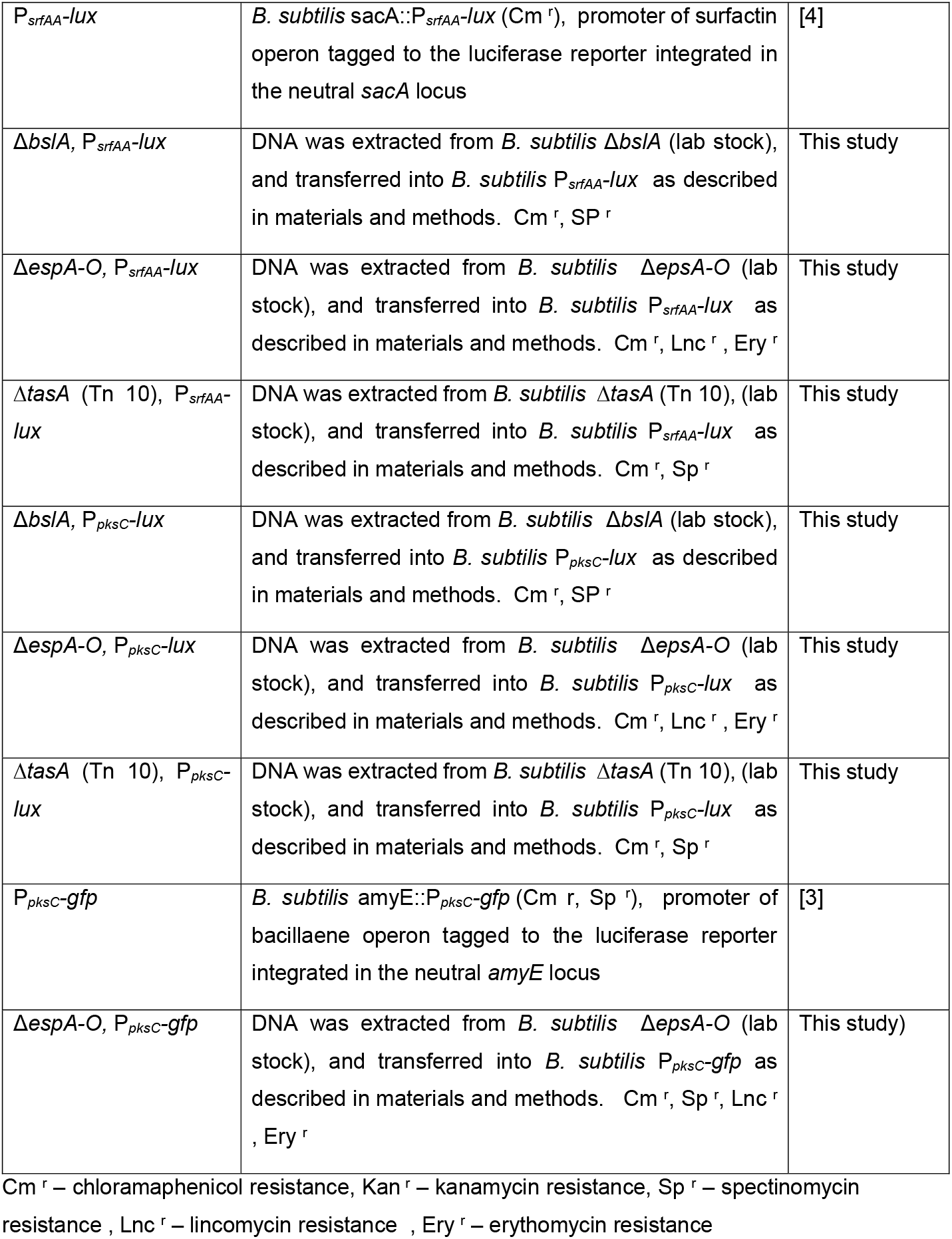
List of strains used in the study.

